# Genetics behind the biosynthesis of nonulosonic acid containing lipooligosaccharides in *Campylobacter coli*

**DOI:** 10.1101/254235

**Authors:** Alejandra Kolehmainen, Mirko Rossi, Jacek Stupak, Jianjun Li, Michel Gilbert, Warren Wakarchuk

**Author notes:** Author for Correspondence: Alejandra Kolehmainen. W.W., M.G. and M.R. contributed equally to this work.

## Abstract

*Campylobacter jejuni* and *Campylobacter coli* are the most common cause of bacterial gastroenteritis in the world. Ganglioside mimicry by *C. jejuni* lipooligosaccharide (LOS) is the triggering factor of Guillain-Barré syndrome (GBS), an acute polyneuropathy. Sialyltransferases from the glycosyltransferase (GT) family 42 are essential for the expression of ganglioside mimics in *C. jejuni*. Recently, two novel GT-42 genes, *cstIV* and *cstV*, have been identified in *C. coli.* Despite being present in ∼11% of currently available *C. coli* genomes, the biological role of *cstIV* and *cstV* is unknown. In the present study, mutation studies in two strains expressing either *cstIV* or *cstV* were performed and mass spectrometry was used to investigate differences in the chemical composition of LOS. Attempts were made to identify donor and acceptor molecules using *in vitro* activity tests with recombinant GT-42 enzymes. Here, we show that CstIV and CstV are involved in *C. coli* LOS biosynthesis. In particular, *cstV* is associated with LOS sialylation, while *cstIV* is linked to the addition of a diacetylated nonulosonic acid residue.

**IMPORTANCE:** Despite being a major foodborne pathogen, *Campylobacter coli* glycobiology has been largely neglected. The genetic makeup of the *C. coli* lipooligosaccharide biosynthesis locus was largely unknown until recently. *C. coli* harbour a large set of genes associated to lipooligosaccharide biosynthesis, including several putative glycosyltransferases involved in the synthesis of sialylated lipooligosaccharide in *Campylobacter jejuni*. In the present study, *C. coli* was found to express lipooligosaccharide structures containing sialic acid and other nonulosonate acids. These findings have a strong impact in understanding *C. coli* ecology, host-pathogen interaction, and pathogenesis.

## INTRODUCTION

Nonulosonic acids are a highly diverse family of nine-carbon α-keto acids. The most naturally abundant nonulosonic acids are the sialic acids (*N-*acetylneuraminic acid, Neu5Ac) and derivatives (1). Initially thought to be only a deuterostome feature, sialic acids have been found in virulence associated bacterial cell surface glycoconjugates such as lipopolysaccharides, capsules, pili, and flagella (2-4). Furthermore, these sialylated structures have been shown to influence pathogenesis through immune evasion, adhesion, and invasion (5, 6). Sialyltransferases catalyse the transfer of sialic acid from cytidine-5′-monophospho-*N*-acetylneuraminic acid (CMP-Neu5Ac) to an acceptor and are key in the synthesis of sialoglycoconjugates. Known sialyltransferases have been classified into seven distinct CAZy (Carbohydrate-active enzymes database) glycosyltransferase (GT) families; GT-29, GT-38, GT-42, GT-52, GT-80, GT-97, and GT-100 (7). In *Campylobacter jejuni,* the most common cause of bacterial gastroenteritis, CMP-Neu5Ac biosynthesis (*neuA, neuB,* and *neuC*) and GT-42 genes are present in the lipooligosaccharide (LOS) biosynthesis locus classes A, B, C, M, R, and V (8-10). *C. jejuni* strains carrying one of these genetic classes synthesize LOS structures generally resembling gangliosides (9, 11-13). In some cases, infection with a *C. jejuni* strain expressing ganglioside-like LOS induces production of cross-reactive anti-ganglioside antibodies. This leads to the development of Guillian-Barré syndrome (GBS); an acute autoimmune polyradiculoneuropathy disease with ∼5% mortality rate (14).

*C. coli*, the second most common cause of campylobacteriosis, has also been isolated from GBS patients (15-18). Nevertheless, the role of *C. coli* in GBS has largely remained unclear due to the seemingly absence of key elements for the synthesis of ganglioside-like LOS (i.e GT-42 and *neuABC* genes). Recently, three newly identified *C. coli* LOS-associated GT-42 genes were reported in the LOS biosynthesis locus: *cstIV, cstV,* and *cstVI* (10, 19, 20). While *cstVI* is generally found as a pseudogene, *cstIV* and *cstV* may potentially be involved in LOS biosynthesis (19). In this manuscript we sought to explore the role of the newly identified GT-42 enzymes CstIV and CstV in *C. coli* LOS biosynthesis.

## RESULTS

### CstIV and CstV are involved in LOS biosynthesis

The LOS of Δ*cstIV* and Δ*cstV* strains showed an increased mobility on silver stained SDS-PAGE gels relative to the WTs (Fig. 1). Thus, deletion of *cstV* in *C. coli* 76339 and *cstIV* in *C. coli* 73 resulted in a truncated LOS. The complemented *cstV* mutant exhibited two LOS bands on SDS-PAGE gels; the upper one corresponding to the WT LOS and the lower molecular weight band to the truncated LOS (Suppl. Fig. S1). This suggests that partial restoration of the phenotype was achieved upon complementation in *cis* of Δ*cstV-*SR4. Deletion of the second putative sialyltransferase of *C. coli* 76339, *cstI* located in the capsular locus, had no effect on LOS mobility on SDS-PAGE gel (data not shown).

**Figure 1.**
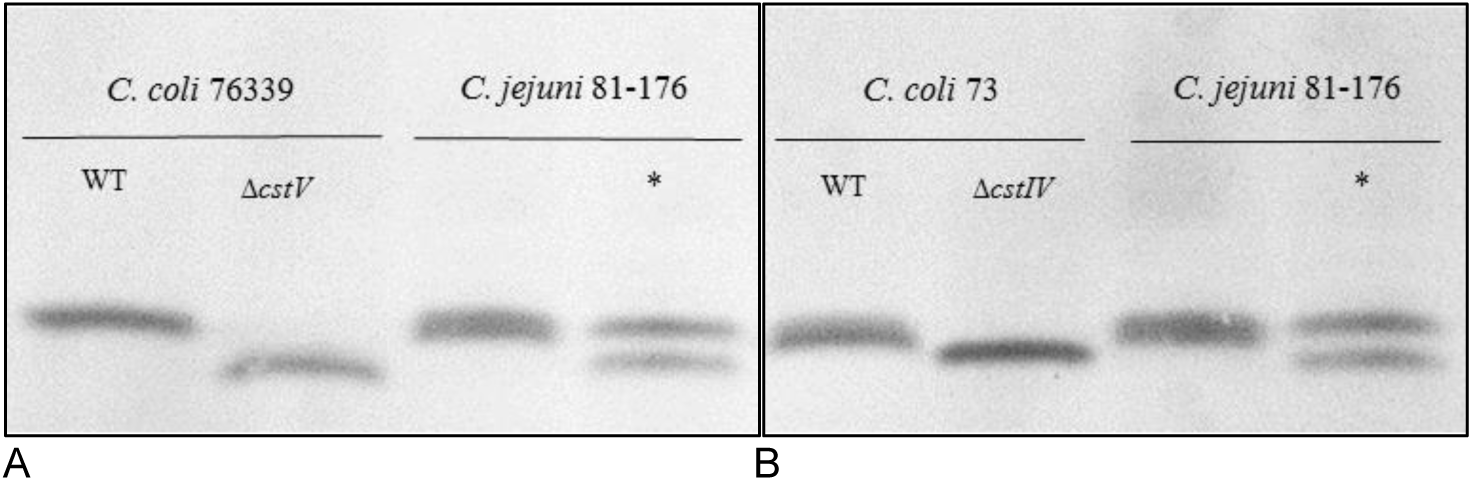
Electrophoresis mobility comparison of (A) *C. coli* 76339 LOS of WT and mutant strains and (B) *C. coli* 73 LOS of WT and mutant strains. *C. jejuni* 81-176 was used as a reference. Lanes marked with an asterisk show *C. jejuni* 81-176 LOS samples treated with neuraminidase to indicate the expected mobility when a Neu5Ac residue is removed.

Though complementation of Δ*cstIV* strain was infeasible, owing to the absence of a suitable loci, it is unlikely that the mobility shift resulting from *cstIV* disruption was due to a polar effect, as *cstIV* is followed by genes that are transcribed in the opposite direction. While *cstIV* and *cstV* were found to be involved in LOS biosynthesis, neuraminidase treatment had no impact on LOS mobility (Suppl. Fig. S2).

### *C. coli* 76339 *neuB1* is involved in the biosynthesis of CstV substrate

Since no clear shift in the electrophoretic mobility of *C. coli* 76339 LOS was detected after neuraminidase treatment (Suppl. Fig. S2), the putative sialic acid synthase, *neuB1,* located downstream from *cstV* (Fig. 2) was knocked out to determine whether CMP-Neu5Ac was the donor molecule for CstV. The LOS of 76339 Δ*neuB1-*SR2 showed a similar profile to those of 76339 Δ*cstV*-SF1 and 76339 Δ*cstV*-SR4 (Suppl. Fig. S3). Thus, inactivation of *neuB1* results in a seemingly similar LOS truncation to the one observed in Δ*cstV* strains, suggesting the potential involvement of *neuB1* in the synthesis of the CstV donor.

**Figure 2.**
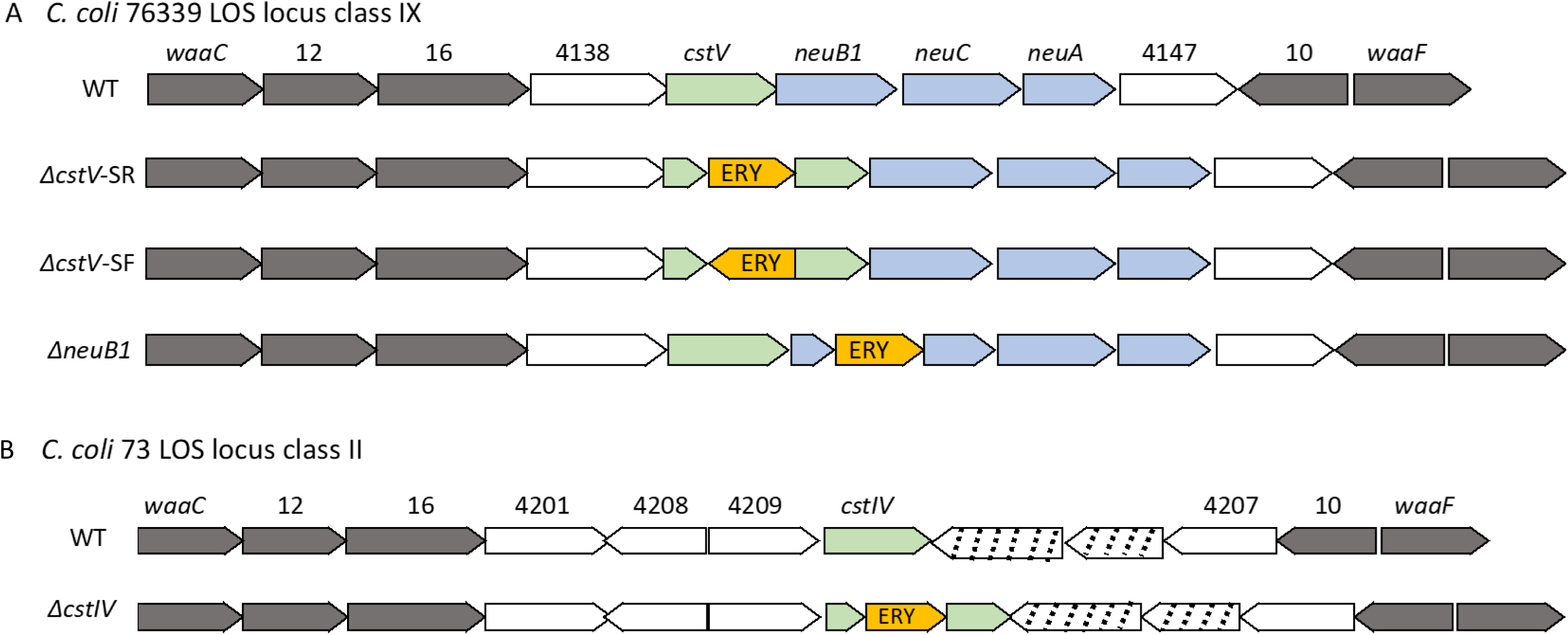
Schematic representation of (A) *C. coli* 76339 LOS locus class IX and (B) *C. coli* 73 LOS locus class II wild types and corresponding mutants. Grey arrows are conserved genes across LOS locus classes as described in (10). Light green arrows represent the GT-42 encoding genes. Light blue arrows represent the genes involved in CMP-Neu5Ac biosynthesis. White arrows show accessory genes with unknown functions. Striped arrows represent pseudogenes in LOS locus class II. Orange arrow indicated with ERY corresponds to the antibiotic resistance cassette used for producing the mutants. Numbers correspond to the gene clusters as described in (19).

*C. coli* 73 lacks Neu5Ac biosynthesis genes orthologues. However, as other *C. coli* strains, it synthetizes other nonulosonic acids through the activity of *neuB2* and *neuB3* genes. Deletion of *neuB2* had no impact on *C. coli* 73 LOS electrophoretic mobility (data not shown) and, despite repetitive attempts, no viable *C. coli* 73 Δ*neuB3* mutants were obtained.

### CstIV and CstV are associated to nonulosonate residues in *C. coli* LOS

Predicted LOS compositions by LC-MS for *C. coli* 76339 WT and mutants are shown on Table 1. *C. coli* 76339 contains a core oligosaccharide linked via two 3-deoxy-D-manno-oct-2-ulosonic acid (Kdo) molecules to a lipid A molecule. The core oligosaccharide of *C. coli* 76339 is composed of heptoses (Hep), hexoses (Hex), hexosamines (HexNAc), and NeuAc. The resulting MS/MS spectrum of *m/z* 1064.0 obtained from the *O*-deacylated LOS of *C. coli* 76339 WT revealed a single ion at *m/z* 1214.4 corresponding to Hex_3_•Hep_2_•PEtn_1_•KDO_1_ (Fig. 3a). The fragment ions at *m/z* 1052.4 and 890.3 correspond to the additional loss of two Hex residues. The spectra also revealed ions that derived from lipid A, m/z 693.5 and *m/z* 388.3 corresponding to HexN3N_1_•P_1_•(C14:0 3-OH)_2_ and HexN3N_1_•(C14:0 3-OH)_1_, respectively. The observation of fragment ions at *m/z* 292.1 and 274.1 provided evidence for the presence of sialic acid on core region LOS. The MS/MS spectrum of precursor ion *m/z* 1064.0 from *C. coli* 76339 Δ*cstI* is similar to that from *C. coli* 76339 WT, in which the diagnostic ions for sialic acid were detected at *m/z* 292.1 and 274.1 (Fig. 3b). However, no sialic acid was detected in the MS/MS spectrum *C. coli* 76339 Δ*cstV* (Fig. 3c). Thus, *cstV* is associated to the presence of NeuAc, while *cstI* plays no role in *C. coli* 76339 LOS biosynthesis.

**Table 1.** LC-MS in positive mode data and proposed compositions for *O*-deacylated LOS of *C. coli* 76339 and corresponding *cstI* and *cstV* knock-out mutants.

| Strain | Observed ions<br>(*m/z*) |  |  |  | Molecular mass<br>(Da) |  | Proposed compositions |  |  |
| --- | --- | --- | --- | --- | --- | --- | --- | --- | --- |
|  | [M+3H]<sup>3+</sup> | [M+2H+NH<sub>4</sub>]<sup>3+</sup> | [M+2H]<sup>2+</sup> | [M+H+NH<sub>4</sub>]<sup>2+</sup> | Observed | Calculated<sup>i</sup> | Core oligosaccharide | Phosphorylation<br>in lipid A | Acylation in<br>lipid A |
| WT | 1063.43 | 1069.11 |  |  | 3187.30 | 3187.35 | Kdo<sub>2</sub>•Hep<sub>2</sub>•Hex<sub>4</sub>•HexNAc<sub>1</sub>•NeuAc<sub>1</sub> | *P*PEtn | 3 *N*-(C14:0 3-OH) |
|  | 1104.43 | 1110.11 |  |  | 3310.30 | 3310.36 | Kdo<sub>2</sub>•Hep<sub>2</sub>•Hex<sub>4</sub>•HexNAc<sub>1</sub>•NeuAc<sub>1</sub>•PEtn<sub>1</sub> | *P*PEtn | 3 *N*-(C14:0 3-OH) |
|  | 1138.48 | 1144.17 |  |  | 3412.46 | 3412.56 | Kdo<sub>2</sub>•Hep<sub>2</sub>•Hex<sub>4</sub>•HexNAc<sub>1</sub>•NeuAc<sub>1</sub> | *P*PEtn | 4 *N*-(C14:0 3-OH) |
|  | 1179.51 | 1185.17 |  |  | 3535.51 | 3535.57 | Kdo<sub>2</sub>•Hep<sub>2</sub>•Hex<sub>4</sub>•HexNAc<sub>1</sub>•NeuAc<sub>1</sub>•PEtn<sub>1</sub> | *P*PEtn | 4 *N*-(C14:0 3-OH) |
| Δ*cstI* | 1063.44 | 1069.11 |  |  | 3187.31 | 3187.35 | Kdo<sub>2</sub>•Hep<sub>2</sub>•Hex<sub>4</sub>•HexNAc<sub>1</sub>•NeuAc<sub>1</sub> | *P*PEtn | 3 *N*-(C14:0 3-OH) |
|  | 1104.44 | 1110.12 |  |  | 3310.33 | 3310.36 | Kdo<sub>2</sub>•Hep<sub>2</sub>•Hex<sub>4</sub>•HexNAc<sub>1</sub>•NeuAc<sub>1</sub>•PEtn<sub>1</sub> | *P*PEtn | 3 *N*-(C14:0 3-OH) |
|  | 1138.51 | 1144.18 |  |  | 3412.52 | 3412.56 | Kdo<sub>2</sub>•Hep<sub>2</sub>•Hex<sub>4</sub>•HexNAc<sub>1</sub>•NeuAc<sub>1</sub> | *P*PEtn | 4 *N*-(C14:0 3-OH) |
|  | 1179.51 | 1185.20 |  |  | 3535.55 | 3535.57 | Kdo<sub>2</sub>•Hep<sub>2</sub>•Hex<sub>4</sub>•HexNAc<sub>1</sub>•NeuAc<sub>1</sub>•PEtn<sub>1</sub> | *P*PEtn | 4 *N*-(C14:0 3-OH) |
| Δ*cstV* |  |  | 1225.56 | 1234.09 | 2449.14 | 2449.14 | Kdo<sub>2</sub>•Hep<sub>2</sub>•Hex<sub>2</sub>•HexNAc<sub>1</sub> | *P* | 3 *N*-(C14:0 3-OH) |
|  |  |  | 1287.08 | 1295.56 | 2572.13 | 2572.15 | Kdo<sub>2</sub>•Hep<sub>2</sub>•Hex<sub>2</sub>•HexNAc<sub>1</sub> | *P*PEtn | 3 *N*-(C14:0 3-OH) |
|  |  |  | 1348.58 |  | 2695.14 | 2695.16 | Kdo<sub>2</sub>•Hep<sub>2</sub>•Hex<sub>2</sub>•HexNAc<sub>1</sub>•PEtn<sub>1</sub> | *P*PEtn | 3 *N*-(C14:0 3-OH) |

<sup>i</sup> Isotope-monoisotopic mass units were used for calculation of molecular mass values based on proposed compositions as follows: HexN, 161.0688; HexN3N, 160.0848; C14:0 3-OH, 226.1933; *PEtn*, 123.0085; *P*, 79.9663; Kdo, 220.0583; Hep, 192.0634; Hex, 162.0528; HexNAc, 203.0794; NeuAc, 291.0954; H<sub>2</sub>O, 18.0106.

**Figure 3.**
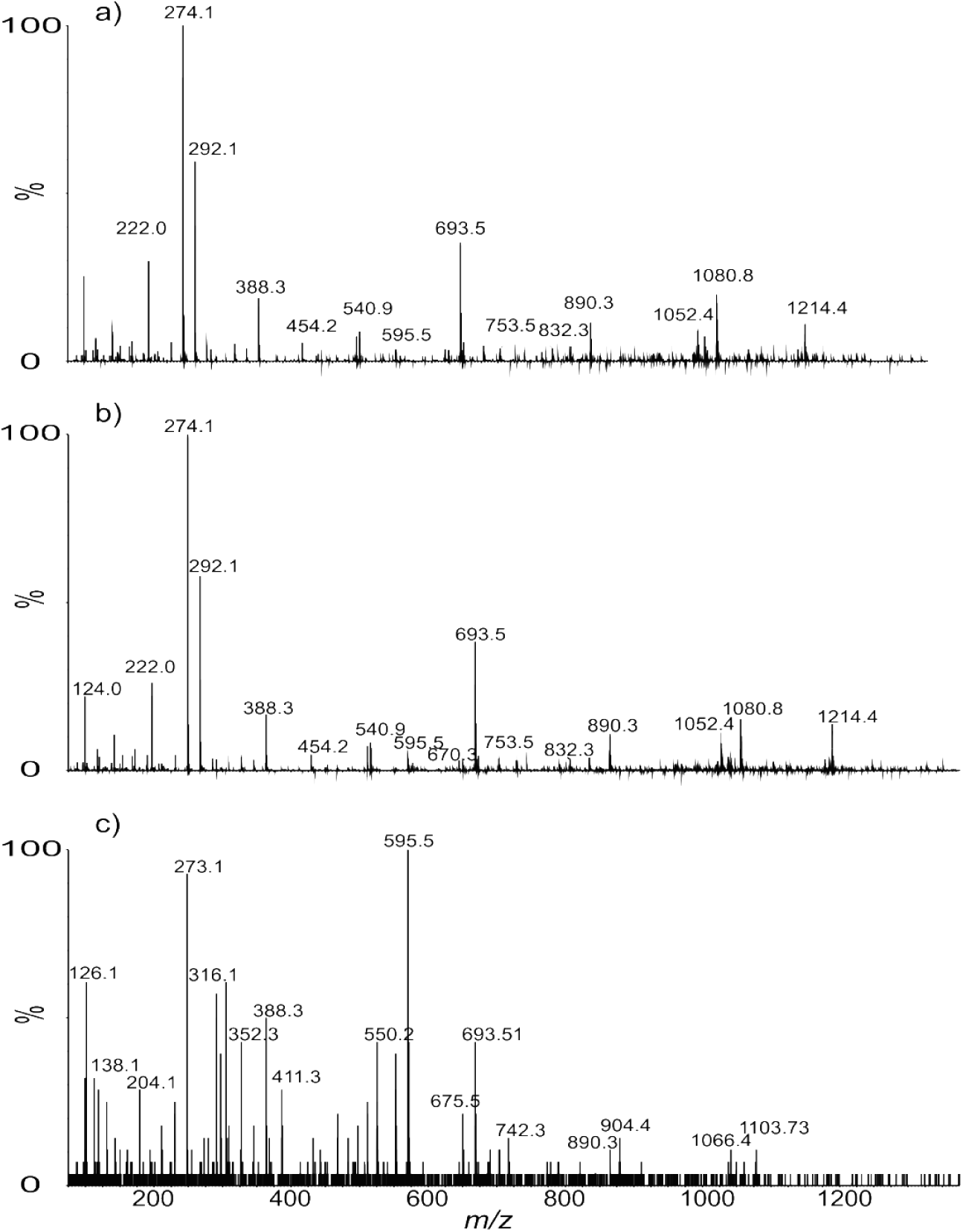
MS/MS spectra for the precursor ions of *O*-deacylated LOS from (a) *C. coli* 76339 WT, *m/z* 1064.0; (b) *C. coli* 76339 Δ*cstI, m/z* 1064.0; and (c) *C. coli* 76339 Δ*cstV, m/z* 1295.4.

A similar lipid A moiety was indicated by the MS/MS spectrum obtained from the *O*-deacylated LOS of *C. coli* strain 73 WT (Fig. 4a). The spectra also revealed ions that derived from lipid A, *m/z* 693.5 and *m/z* 388.3 corresponding to HexN3N_1_•P_1_•(C14:0 3-OH)_2_ and HexN3N_1_•(C14:0 3-OH)_1_, respectively. The observation of fragment ions at *m/z* 317.2 and 299.1 provided evidence for the presence of a residue with a molecular weight of 334.2 Da or 316.2 Da for its anhydrous form on core region LOS. These masses are consistent with free diNAc-nonulosonate and its conjugated form, respectively. However, these characteristic ions were not detected in the MS/MS spectrum *C. coli* 73 Δ*cstIV*-SF3 (Fig. 4b). Thus, suggesting the role of *cstIV* in the biosynthesis of diNAc-nonulosonate LOS in *C. coli* 73.

**Figure 4.**
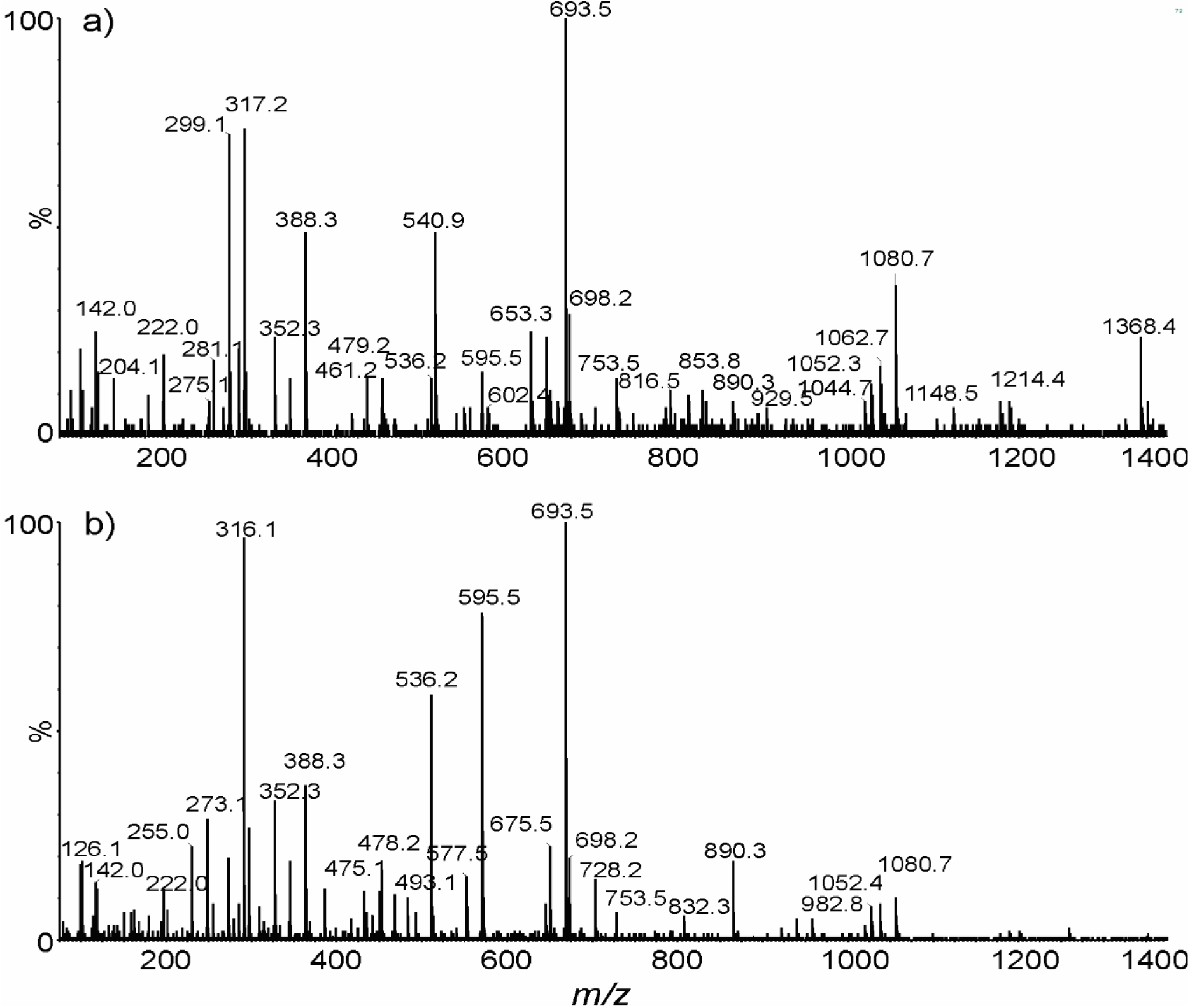
MS/MS spectra for the precursor ions of *O*-deacylated LOS from (a) *C. coli* strain 73 WT, *m/z* 1072.4; (b) *C. coli* 73 Δ*cstIV*-SF3, *m/z* 1566.0.

### No sialyltransferase activity was detected for CstIV and CstV using *in vitro* assays

To determine whether CstIV and CstV are capable of transferring Neu5Ac, *C. coli* crude protein extracts were tested for sialyltransferase activity using sugar acceptors labelled with either boron-dipyrromethene (abbreviated as BODIPY or BDP) or fluorescein (FCHASE). No sialyltransferase activity was detected in the crude protein extracts of *C. coli* 73 and *C. coli* 73 Δ*cstIV*-SF3. Monospecific α-2,3-sialyltransferase activity was detected in *C. coli* 76339 WT, *C. coli* 76339 Δ*cstV*-SF1, and *C. coli* 76339 Δ*cstV*-SR4 crude protein extracts using BDP-Lactose (BDP-Lac) and BDP-*N*-acetyllactosamine (BDP-LacNAc) (Suppl. Fig. S4). Since the genome of *C. coli* 76339 is known to carry the gene encoding the monospecific CstI α-2,3-sialyltransferase, the assays were also performed using *C. coli* 76339Δ*cstI-*XR3 and *C. coli* 76339Δ*cstV*-SRΔ*cstI*-XR1 protein extracts (data not shown). No measurable enzymatic activity was detected with any of the tested acceptors in these Δ*cstI* strains which demonstrated that the activity detected in *C. coli* 76339 WT was due to CstI. Furthermore, all tested recombinant CstIV and CstV showed no activity with any of the tested acceptors, suggesting that either none of the tested glycans was a suitable acceptor or that another nonulosonate is the actual donor for these enzymes (data not shown).

## DISCUSSION

*C. jejuni* GT-42 were the first glycosyltransferases from this CAZy family to be enzymatically and structurally characterized; CstII variants can be either monofunctional α2,3-sialyltransferases or bifunctional α2,3-/α2,8-sialyltransferase, while CstI and CstIII are monofunctional α2,3-sialyltransferases (21-24). CstII and CstIII activity has been shown to be essential for the biosynthesis of ganglioside-like LOS structures, which are linked to GBS onset (12, 24). Despite the importance of GT-42 enzymes in virulence and pathogenesis (25-28), the activity of these glycosyltransferases has not been explored in other *Campylobacter* species. Approximately 29% of *C. coli* genomes have been found to contain a GT-42 encoding gene within the LOS biosynthesis locus (19). While *cstVI* was the most common LOS associated GT-42 encoding gene in *C. coli*, in 99% of the analysed genomes it was observed to be present as a pseudo gene (19). Thus, we focused our attention on the role of *cstIV* and *cstV* in LOS biosynthesis. Until recently, *cstV* had been solely identified in the genome of *C. coli* 76339 (20). However, in a systematic screen of publicly available *C. coli* genomes several *cstV* positive strains were identified (19). Since *in vitro* assays have been previously used to determine the activity of *C. jejuni* GT-42 enzymes (9, 21), a similar approach was attempted to define CstIV and CstV activity. *C. coli* 76339 crude protein extracts were tested for sialyltransferase activity, as Neu5Ac had been previously detected in the strain’s LOS (20). Monofunctional sialyltransferase activity was initially observed but was found to be due to CstI activity. As in *C. jejuni, C. coli* 76339 *cstI* is located outside the LOS biosynthesis locus, and encodes an α2,3-sialyltransferase which has no role in LOS biosynthesis (20, 21). Although transcriptomic analysis showed polycistronic expression of LOS biosynthesis genes, indicating the active expression of *cstV* (data not shown), no sialyltransferase activity was detected on the protein extracts of the *cstI* mutant strain. Inactivation of *neuB1* or *cstV* resulted in identical LOS electrophoretic profiles. Additionally, LC-MS analysis showed that the inactivation of *cstV* resulted in the loss 2 Hex and 1 NeuAc residues. Nevertheless, recombinant CstV exhibited no detectable activity with any of the tested acceptors. Thus, it is very likely that *cstV* is associated to *C. coli* 76339 LOS sialylation. Yet, further studies are required to identified CstV natural acceptor and corroborate its activity *in vitro.*

After *cstVI, cstIV* is the most common orthologue; being present in ∼38% of the genomes positive for a LOS associated GT-42. Previously, no evidence of Neu5Ac had been found in the LOS of strains containing a *cstIV* orthologue (29). This was to be expected as Neu5Ac biosynthesis genes are rarely present in strains carrying *cstIV* (19). Furthermore, no sialyltransferase activity was detected neither in *C. coli* 73 protein extracts nor in recombinant CstIV. Nevertheless, deletion of *cstIV* in *C. coli* 73 resulted in a truncated LOS. Thus, suggesting a link between *cstIV* and LOS biosynthesis. Sequence alignment of CstIV with previously characterized GT-42 sialyltransferases revealed numerous amino acid substitutions at conserved positions (Suppl. Fig. S5) (30). Additionally, superimposition of CstIV on *C. jejuni* CstII structure identified various substitutions at amino acids involved in substrate interactions (23, 31-33). Interestingly, most substitutions predicted to impact CstIV activity were in the amino acids associated with CMP-Neu5Ac, particularly with the Neu5Ac moiety. Moreover, these substitutions were conserved in multiple CstIV orthologues (23, 32, 33). Altogether, results pointed at the possibility of an alternative sugar donor for CstIV. Detection of a diNAc-nonulosonate residue in *C. coli* 73 WT LOS and its absence in *C. coli* 73 Δ*cstIV*-SF3 prompted an investigation on genes potentially linked to the synthesis of this residue. In *C. coli, neuB2* (*ptmC, legI*) and *neuB3* (*pseI*) are conserved flagella glycosylation genes involved in the synthesis of legionaminic and pseudaminic acid derivatives, respectively (34-40). Deletion of *neuB2* had no impact on *C. coli* 73 LOS electrophoretic mobility, implying that *neuB2* is not involved in the synthesis of CstIV donor. Despite repetitive attempts, no viable *C. coli* 73 Δ*neuB3* mutants were obtained. Although *neuB3* deletion has been successful in *C. coli* VC167, disruption of flagellin glycosylation and the potential truncation of the LOS might have resulted in a lethal phenotype for *C. coli* 73 (40). In sum, it is tempting to speculate that the diNAc-nonulosonate residue in *C. coli* 73 WT corresponds to pseudaminic acid. However, the nature of this residue cannot be inferred from MS/MS spectra alone since many diNAc-nonulosonate variants have been identified (41).

In conclusion, although we could not determine the complete structures of the LOS outer cores of *C. coli* 73 and *C. coli* 76339, we have established that they both contain nonulosonic acid. We have also unequivocally demonstrated that CstIV and CstV are involved in the synthesis of LOS in their respective stains and, more specifically, they are responsible of the transfer of a nonulosonic acid residue to the outer core.

## METHODS

### Bacterial strains, plasmids, and growth conditions

Bacterial strains used in this study are listed in Table 2. Two *C. coli* strains, expressing either *cstIV* or *cstV*, were selected. Figure 2 shows a schematic representation of the LOS locus and the position of the insertion of the antibiotic resistance cassette for the mutational studies. *C. coli* 73 possesses LOS locus class II, which contains a copy of *cstIV* and lacks sialic acid biosynthesis genes *neuABC* or other copies of putative sialyltransferases. However, strain 73 possesses the conserved biosynthesis pathways for both legionaminic (including legionaminic acid synthetase *neuB2*) and pseudaminic acid (including pseudaminic acid synthetase *neuB3*) in the flagella glycosylation region (29). *C. coli* 76339 harbours two copies of putative sialyltransferases: *cstV* as part of LOS locus class IX (Fig. 2) and a *C. jejuni cstI* orthologue located in the capsule region (19, 20). In addition to the conserved legionaminic and pseudaminic acid biosynthesis pathways, as in *C. coli* 73, *C. coli* 76339 possesses all the genes for the biosynthesis of sialic acid (*neuAB1C*) within the LOS locus class IX (Fig. 2). *C. coli* cultivation and DNA isolation were carried out as previously described, unless specified otherwise (20).

**Table 2.** Bacterial strains

| Strain or plasmid | Genotype and/or phenotype | Reference or source |
| --- | --- | --- |
| *C. coli* 76339 | *cstI*, *cstV*, *neuB* | (20) |
| *C. coli* 76339 Δ*cstV*-SF1 | *cstI*, Δ*cstV*: Ery, *neuB* | This study |
| *C. coli* 76339 Δ*cstV*-SR4 | *cstI*, Δ*cstV*: Ery, *neuB* | This study |
| *C. coli* 76339 Δ*neuB*-SR1 | *cstI*, Δ*neuB*: Ery, *cstV* | This study |
| *C. coli* 76339 Δ*cstI*-XR3 | Δ*cstI*: CAT, *cstV*, *neuB* | This study |
| *C. coli* 76339 Δ*cstV*-SR4 Δ*cstI*-XR1 | Δ*cstI*: CAT, Δ*cstV*: Ery, *neuB* | This study |
| *C. coli* 76339 Δ*neuB*-SR2 | *cstI*, Δ*neuB*: Ery, *cstV* | This study |
| *C. coli* 76339 Δ*cstV*-SR4 Δggt:*cstV*-2 | *cstI*, Δ*cstV*: Ery, *neuB*,<br>Δggt:*cstV*:CAT | This study |
| *C. coli* 73 | *cstIV* | (29) |
| *C. coli* 73 Δ*cstIV*-SF3 | Δ*cstIV*:Ery | This study |
| *C. coli* 73 Δ*neuB*2 | Δ*neuB*2:Ery | This study |
| *C. jejuni* 81-176 |  |  |
| *E. coli* AD202 |  | (46) |

### Construction of Δ*cstIV*, Δ*cstV,* Δ*cstI,* and Δ*neuB* mutants

Chromosomal mutant strains of *C. coli* 76339 (19, 20) and *C. coli* 73 (19, 29) were generated by homologous recombination with suicide vectors containing genes inactivated by the insertion of an antibiotic resistance cassette. All recombinant plasmids and primers are shown in Supplemental material (Suppl. Fig. S6-S11). The genes *cstIV, cstV, neuB1, neuB2,* and *neuB3* were inactivated by the insertion of an erythromycin resistance cassette (EryC) (42), while *cstI* was disrupted with a chloramphenicol acetyltransferase cassette (CAT) (43). The inactivation of *cstV* in *C. coli* 76339 was performed by inserting *eryC* cassette either in the direction of the gene (SR) or in the opposite direction (SF). Preparation of electrocompetent cells and transformation was done as previously described (43). Selection of the mutants was done on nutrient blood agar (NBA) supplemented containing either 10 μg ml^−1^ of erythromycin or 12.5 μg ml^−1^ of chloramphenicol. Homologous recombination of all mutants was verified by PCR. Figure 2 shows a summary of the mutations performed in the LOS locus of *C. coli* strains 76339 and 73.

### Complementation studies

Complementation of *C. coli* 76339 Δ*cstV-*SR4 was done in *cis* by integration of *cstV* under the active promoter of gamma glutamyltranspeptidase (*ggt*). The *ggt* is an accessory gene in *C. coli* and has no role in LOS biosynthesis. Additionally, the *ggt* locus is located far from the LOS locus and its deletion does not induce a loss in bacterial viability. The suicide vector containing an inactivated *ggt* by the insertion of a *cstV* and CAT (pGEM-ggt-cstV-CAT) (Suppl. Fig. S12) was used to transform *C. coli* 76339 Δ*cstV-*SR4 electrocompetent cells as above. Transformants were selected on NBA supplemented with 12.5 μg ml^−1^ of chloramphenicol. Homologous recombination of mutants was verified by testing for GGT activity as before (44). Complementation of *cstIV* was not possible due to the absence of a suitable locus.

### LOS silver staining

LOS profiles were assessed by silver staining as described earlier (29). Additionally, LOS sensitivity to neuraminidase was assessed by treating crude LOS with 2 IU/ml of *Clostridium perfringens* neuraminidase (Sigma-Aldrich) overnight at 37 °C.

### Mass spectrometry analysis of *C. coli* LOS composition

Following 1% formaldehyde in PBS (pH 7.4) treatment, *C. coli* cell pellets were washed 3X in PBS and lyophilized. Then, cells were dehydrated by a sequence of 2 washes in each of 70% ethanol (in PBS), 100% ethanol, and 100% acetone. The dehydrated cells were treated with proteinase K, RNAse A, and DNAse I as previously described (45). Digested cells were then treated with hydrazine to cleave *O*-linked fatty acids (45). The *O*-deacylated LOS samples were analysed by LC-MS by coupling a Waters Premier Q-TOF with an Agilent 1260 capillary LC system. Mass spectrometry was operated in positive-ion detection mode. Liquid chromatography separation was done on an Agilent Eclipse XDB C8 column (5 μm, 50 x 1 mm). The flow rate was 20 μL/min. Solvent A: aqueous 0.2% formic acid/0.028% ammonia; solvent B: Isopropanol with 0.2% formic acid/0.028% ammonia. The following gradient was used: 0-2 min. 10% B, 2-16 min linear gradient to 85% B, 16-25 min. 85% B, 25-30 min., and equilibration at 10% B.

### Sialyltransferase activity test in *C. coli* protein extracts

To test for sialyltransferase activity, *C. coli* 76339 and 73 were grown for 16 h in nutrient broth 2 (Oxoid) (100 rpm, microaerobic atmosphere, and 37 °C). Cells were harvested by centrifugation (10,000 × *g* for 15 min at 4°C) and resuspended in 50 mM HEPES pH 7.5 containing a protease inhibitor cocktail (Sigma). Cells were then lysed by mechanical disruption and debris was removed by centrifugation (10,000 × *g* for 15 min at 4°C). Sialyltransferase activity of protein extracts was tested on boron-dipyrromethene or BODIPY (BDP) labelled Lac, LacNAc, and 3’Sialyllactose or fluorescein (FCHASE) labelled α-GalNAc, β-GalNAc, GM3, α-Gal, β-GlcNAc, α-Glc, β-Glc, Hep-Hep-Glc, as acceptors. Reactions were performed at 37°C in 10 μl volumes containing 50 mM HEPES pH 7.5, 10 mM MgCl_2_, 1 mM CMP-NeuAc, 0.5 mM labelled acceptor, and 6 μl of extract. To stop enzymatic reactions an equal volume of 80% acetonitrile was added. Enzymatic activity was assessed by thin-layer chromatography on silica using a solvent system of ethyl acetate/methanol/water/acetic acid 4:2:1:0.1.

### Expression and activity of recombinant *C. coli* GT-42 enzymes

Gene *cstIV* from *C. coli* 73 and *cstV* from *C. coli* 76339 were amplified and ligated to pCW and pCW-MalET plasmids (46). Ligation products were then electroporated in *E. coli* 10β for plasmid amplification. After sequence confirmation, plasmids were electroporated into *E. coli* AD202 or BL21 for protein expression. Cells containing the protein expression vectors were grown in 200 mL of 2YT medium supplemented with 150 μg/mL ampicillin and 0.2% of glucose at 25°C with 250 rpm shaking. After reaching an A_600nm_ of ∼0.6, protein over-expression was induced with 0.5 mM of isopropyl-β-D thiogalactopyranoside (IPTG) and cultures were further incubated for 16 h. Cells were harvested by centrifugation (10,000 × *g* for 15 min at 4°C) and crude protein extracts were run in 12% SDS-PAGE gels and stained with Coomassie blue to verify overexpression. In addition, *cstIV* and *cstV* genes were synthesized with a T7 promoter and a ribosome binding site upstream of the coding sequence and a T7 terminator downstream from the stop codon (Thermo Scientific). Synthesized products were inserted into pMA-T vector backbone and proteins were synthesized using the cell-free PURExpress *in vitro* Protein Synthesis Kit (New England Biolabs Inc.). Recombinant proteins were screened for sialyltransferase activity as described above.

## Supporting information

## ACKNOWLEDGEMENTS

This research project was supported by the University of Helsinki research grant n. 313/51/2013, and the Walter Ehrström Foundation travel grant. A. K was supported by the Microbiology and Biotechnology graduate program from the University of Helsinki. The authors wish to thank Marja-Liisa Hänninen for providing the strains and Arnoud HM van Vliet for providing the erythromycin resistance cassette. We thank Denis Brochu for help with the preparation of the samples for mass spectrometry analysis and data presentation.

## CONTRIBUTIONS

A.K and M.R designed and coordinated the study. A.K generated all *C. coli* mutants. A.K, M.G, and W.W participated in enzymatic assays. J.S and J.L performed LC-MS analysis. J.S, J.L, and M.G interpreted LC-MS data. A.K drafted the manuscript. All authors have contributed to data interpretation, have critically reviewed the manuscript, and approved the final version as submitted.

## ADDITIONAL INFORMATION

The authors declare that they have no competing interests.

## DISCLAIMER

M. R is currently employed with the European Food Safety Authority (EFSA) in its BIOCONTAM Unit that provides scientific and administrative support to EFSA’s scientific activities in the area of Microbial Risk Assessment. The positions and opinions presented in this article are those of the authors alone and are not intended to represent the views or scientific works of EFSA.

