## Supplementary material for "Genetics behind the biosynthesis of nonulosonic acid containing lipooligosaccharides in *Campylobacter coli*"

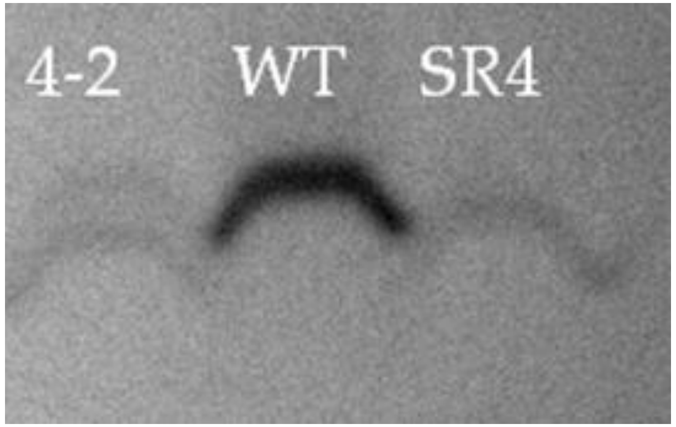


**Supplemental Figure S1.** Electrophoresis mobility comparison of *C. coli* 76339Δ*cstV*Δ*ggt*:*cstV* (4-2), *C. coli* 76339 WT, and *C. coli* 76339Δ*cstV* (SR4).

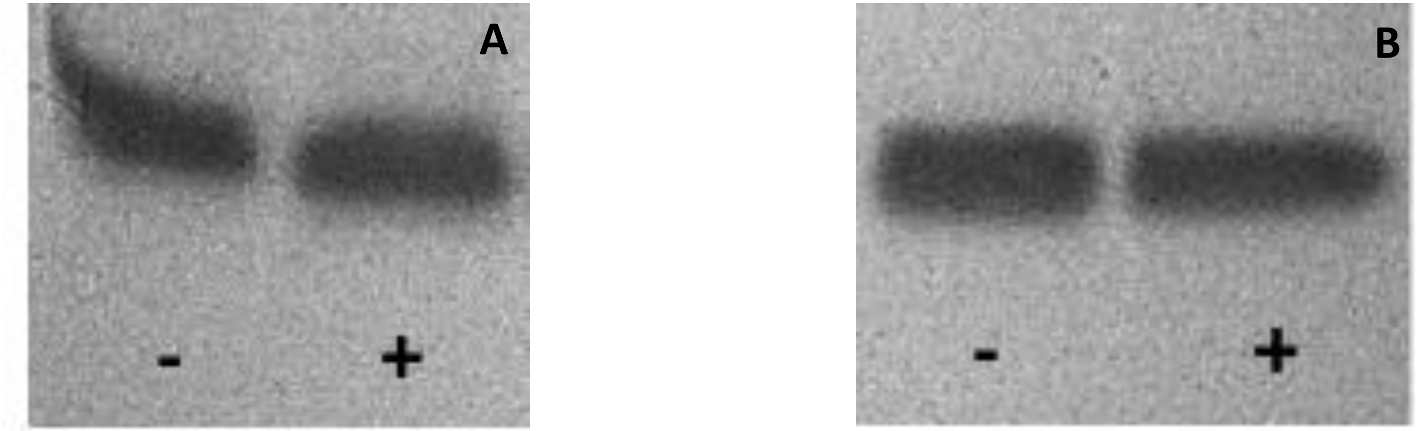


**Supplemental Figure S2.** Electrophoresis mobility comparison of neuraminidase treated(+) and untreated(-) crude LOS. (A) *C. coli* 76339 WT (*cstV*); (B) *C. coli* 73 WT (*cstIV*).

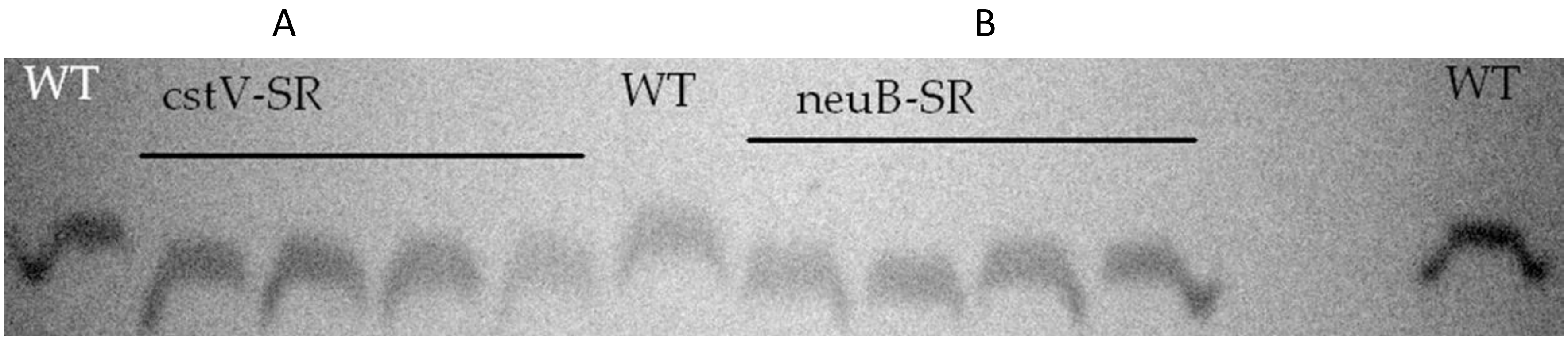


**Supplemental Figure S3.** Electrophoresis mobility comparison of *C. coli* 76339 WT, Δ*cstV,* and Δ*neuB1.* (A) *C. coli* 76339 WT, Δ*cstV*-SR (erythromycin resistance cassette cloned in same direction as *cstV*). (B) Δ*neuB*-SR (erythromycin resistance cassette cloned in same direction as *neuB1*).

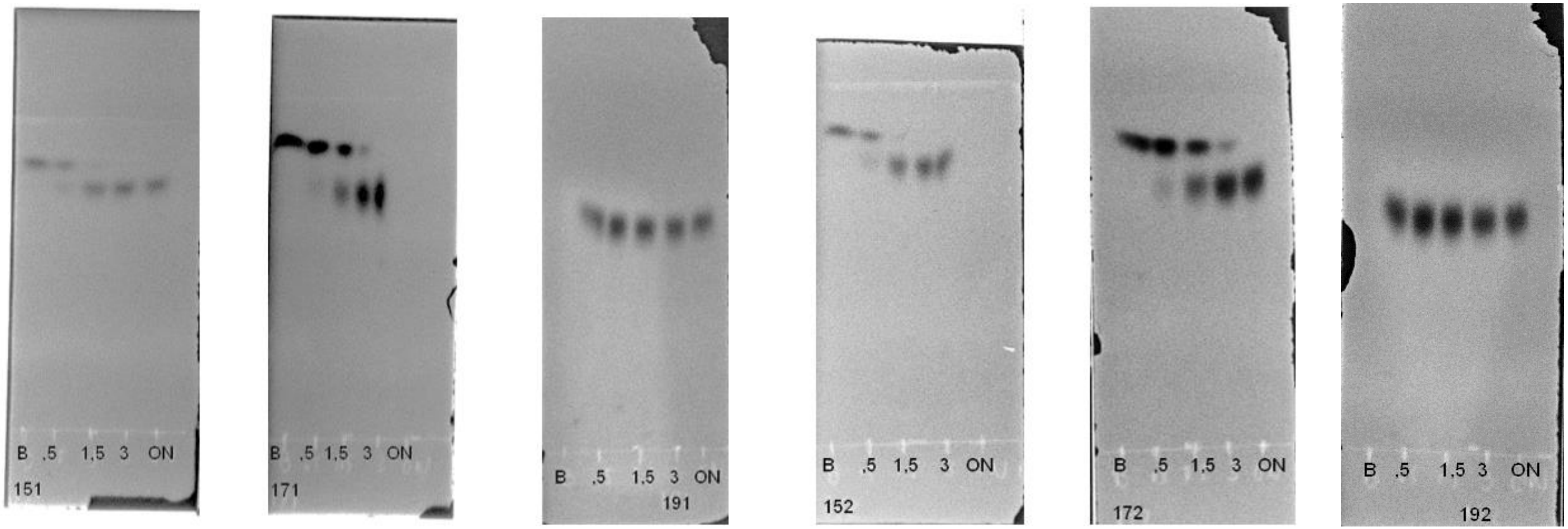


| **Sample code** | **Strain** | **BDP-acceptor** |
|---|---|---|
| **151** | *C. coli* 76339 | Lac |
| **171** | *C. coli* 76339 | LacNAc |
| **191** | *C. coli* 76339 | 3’Sialyllactose |
| **152** | *C. coli* 76339 Δ*cstV* | Lac |
| **172** | *C. coli* 76339 Δ*cstV* | LacNAc |
| **192** | *C. coli* 76339 Δ*cstV* | 3’Sialyllactose |

**Supplemental Figure S4.** Example of enzymatic test. Sialyltransferase activity of protein extracts was tested on BODIPY labelled Lac, LacNAc, and 3’Sialyllactose as donors. Reactions were performed at 37 °C in 10 μL volumes containing 50 mM HEPES pH 7.5, 10 mM $MgCl_2$, 1 mM CMP-NeuAc, 0.5 mM labelled acceptor, and 6 μL of extract. Enzymatic activity was assessed by thin-layer chromatography on silica using a solvent system of ethyl acetate/methanol/water/acetic acid 4:2:1:0.1.
Samples were incubated for 0.5, 1, 1.5, and 3 hours, and overnight (ON)

```
Bi-Cj-cstII ---------------MKKVIIAGNGPSLKEIDYSRLPNDFDVFRCNQFYFEDKYYLGKKCKAVFYNPILFFEQYYTLKHL 65
Cj-cstI     MTRTRMENELIVSKNMQNIIIAGNGPSLKNINYKRLPREYDVFRCNQFYFEDKYYLGKKIKAVFFNPGVFLQQYHTAKQL 80
cstIV       -------MNKNLNFTKKSVLIAGNGPSLKEIDYSLLPKEYDIFRCNHFYAEDKYYTGKKIKAAFFVIPYFFNEYYTMQKM 73
cstV        ------------MIENNAVVVAGNGTSLKEIDYSRLPKEFDVFRCNQFYFEDKYYLGRKVKAAFSFPGVFFEQYYTLNTL 68
HAC1267     -------------MNKKPLIIAGNGPSIKDLDYALFPKDFDVFRCNQFYFEDKYYLGREIKGVFFNAHVFDLQMKITKAI 67
HAC1268     -----------MGTIKKPLIIAGNGPSIKDLDYALFPKDFDVFRCNQFYFEDKYYLGREIKGVFFNPCVLSSQMQTVQYL 69
HBS-02      -------------MPLKPLIVAGNGPSIKDLDYSLFPPDFDVFRCNQFYFEDKYYLGKEVKGVFFNAQVFDMQMKTAREL 67
Pm70        ------MDKFAEHEIPKAVIVAGNGESLSQIDYRLLPKNYDVFRCNQFYFEERYFLGNKIKAVFFTPGVFLEQYYTLYHL 74

Bi-Cj-cstII IQNQEYETELIMCSNYNQAHLENEN-----FVKTFYDYFPDAHLGY-DFFKQLKDFNAYFKFHEIYFNQ--RITSGVYMC 137
Cj-cstI     ILKNEYEIKNIFCSTFNLPFIESND-----FLHQFYNFFPDAKLGY-EVIENLKEFYAYIKYNEIYFNK--RITSGVYMC 152
cstIV       LQNGDYECENIVCKMYNFQDRKE-----KIFRENFKYFFPAAINGYDAFFYKLKELSNMIDFDFCYEYNTTEMLTGTYVI 148
cstV        MQNKEYYCENIVCKLFPLQHEINQKSL-RNFKKIFPLFFPYALDGNEHYFNKLKELNSFINFNFLYDEG-LQITTGMYAI 146
HAC1267     VKNGEYHPDHIYCTHVEPYGYVNGNQQ---LMQEYLEKHFVGVRSTYAYLKDLEPFFILHSKYRNFYDQ--HFTTGIMML 142
HAC1268     MDNGEYSIERFFCSVSTDRHDFDGDYQTILPVDGYLKAHYPFVCDTFSLFKGHEEILKHVKYHLKTYSK--ELSAGVLML 147
HBS-02      SLRQEYYFEDLFCSTIAPFMNFGNHYT---HAQDYLDKHYPGARNTYALLQSLEPFYKLYTTRRNFYQQ--HFTTGVMMI 142
Pm70        KRNNEYFVDNVILSSFNHPTVDLEK------SQKIQALFIDVINGYEKYLSKLTAFDVYLRYKELYENQ--RITSGVYMC 146

Bi-Cj-cstII AVAIALGYKEIYLSGIDFYQN-GSSYAFDTKQKNLLKLAPNFKNDNSHYIGHSKNTDIKALEFLEKTYKIKLYCLCPNSL 216
Cj-cstI     AIAIALGYKTIYLCGIDFYEG-DVIYPFEAMSTNIKTIFPGIK-DFKPSNCHSKEYDIEALKLIKSIYKVNIYALCDDSI 230
cstIV       CCAVACGYKEIYLAGMDFGDE-KYNYFSEKIEK-----IKESKKR--TTKMHHKSIDLKILDFLQQQYNVKIFSICPNSS 220
cstV        ACAVACGYKEIYITGIDFYST-QEYAFDIKDKIGLYALNPSFKIQ--YLKSHSKETDLEIILSFLKQTYNANFFSISPKSP 223
HAC1267     LVAIQLGYKEIYLCGIDFYENGFGHFYE----------NQGGFFEEDSDPMHDKNIDIQALELAKKY--AKIYALVPNSA 210
HAC1268     LSAVVLGYKEIYLVGIDFGASSWGHFYDE---------SQSQHFSNHMADCHNIYYDMLTICLCQKY--AKLYALAPNSP 216
HBS-02      IVAIVLGYKEIYCAGLDFYLEGLGHFYH----------VKSPHFTLAPDCQHTKDLDIKGIEVAKQY--AQIYALVPNSA 210
Pm70        AVAIAMGYTDIYLTGIDFYQASEENYAFDNKKPNIIRLLPDFRKEKTLFSYHSKDIDLEALSFLQQHYHVNFYSISPMSP 226

Bi-Cj-cstII LANFIELAPN---------------------------------------------------------------------- 226
Cj-cstI     LANHFPLSIN---------------------------------------------------------------------- 240
cstIV       INAFIPLHPI---------------------------------------------------------------------- 230
cstV        MTKYIPLAPK---------------------------------------------------------------------- 233
HAC1267     LVKMIPLSSQKGVLEKVKDRIGLGEFKREKFGQKELERQKELERQKELERQKELERQKELERQKELERQKELERQKELER 290
HAC1268     LSHLLTLNPQAKYPFELLDKP-IGYTSDLIISSPLEEKLLEFKNIEEKLLEFKNIEEKLLEFKNIEEKLLEFKNIEEKLL 295
HBS-02      LSAILPLSPHKNALSQEK-------------------------------------------------------------- 228
Pm70        LSKHFPIPTV---------------------------------------------------------------------- 236

Bi-Cj-cstII -------------------------LNSN--FIIQEKNN-YTKDILIPSSEAYGKFSKNINF------------------ 260
Cj-cstI     -------------------------INNN--FTLENKHNNSINDILLTDNTPGVSFYKNQLKAD----NKIMLNFYNILH 289
cstIV       -------------------------QNNENIFKPIERPKDAIKTQLTPPIKAVRRYKR-LYLES----NIIIKFFHELVQ 280
cstV        -------------------------QNYS--FDIEEKSSESIKDFLIPSKKAYRNYSRALYLQN----NMFYNFIHDCLK 282
HAC1267     QKELERQKELERQKELERQKELERQKELERQKELERQKELERQKELELERSLKARLKAVLASKGIRGDNLIIVSLKDTYR 370
HAC1268     EFKNIEEKLLEFKNIEEKLLEFKNIEEKLLEFKNIEEKLLEFKNIEEKLLEFKNIEEKLLEFKNIEEKLLEFKNIEEKLL 375
HBS-02      -------------------------MQALKLGDPKPNGYIDDVCVDPVEFSMRATLIQAIQSVGITPDNLIYKGLNMVWR 283
Pm70        -------------------------EDDCETTFVAPLKENYINDILLPPHFVYEKLGTIVSKKSRFHSNLIVRLIRDLLK 291

Bi-Cj-cstII ----------------------------------------------- 260
Cj-cstI     SKDNLIKFLNK------------------------------------ 300
cstIV       VPRRIRHYYSKTRYTR------------------------------- 296
cstV        FPSALKNYFKNIKK--------------------------------- 296
HAC1267     LFKGGFALLLD------LKALKSIIKAFLKR---------------- 395
HAC1268     EFKNIEEKLLASRLNNILRKIKRKILPFFWGGGVTPTLKVSFRWGAA 422
HBS-02      CASDLYRVVRG------LYRLSLKALYFLRAWFKRHRATGG------ 318
Pm70        LPSALKHYLKEK----------------------------------- 303
```

| **Important residues for CMP** | |
|---|---|
| **CstII** | **CstIV** |
| Thr131 | Leu142 |
| Ser132 | Thr143 |
| **Important residues for Neu5Ac** | |
| **CstII** | **CstIV** |
| Gln32 | His40 |
| Asn51 | Val59 |
| Gln58 | Glu66 |
| Arg129 | Glu140 |
| Ser132 | Thr143 |
| Tyr185 | Thr189 |

**Supplemental Figure S5.** Multiple sequence alignment of characterized GT-42 sialyltransferases and *C. coli* 76339 *cstV* and *C.coli* 73 *cstIV.*

**A**

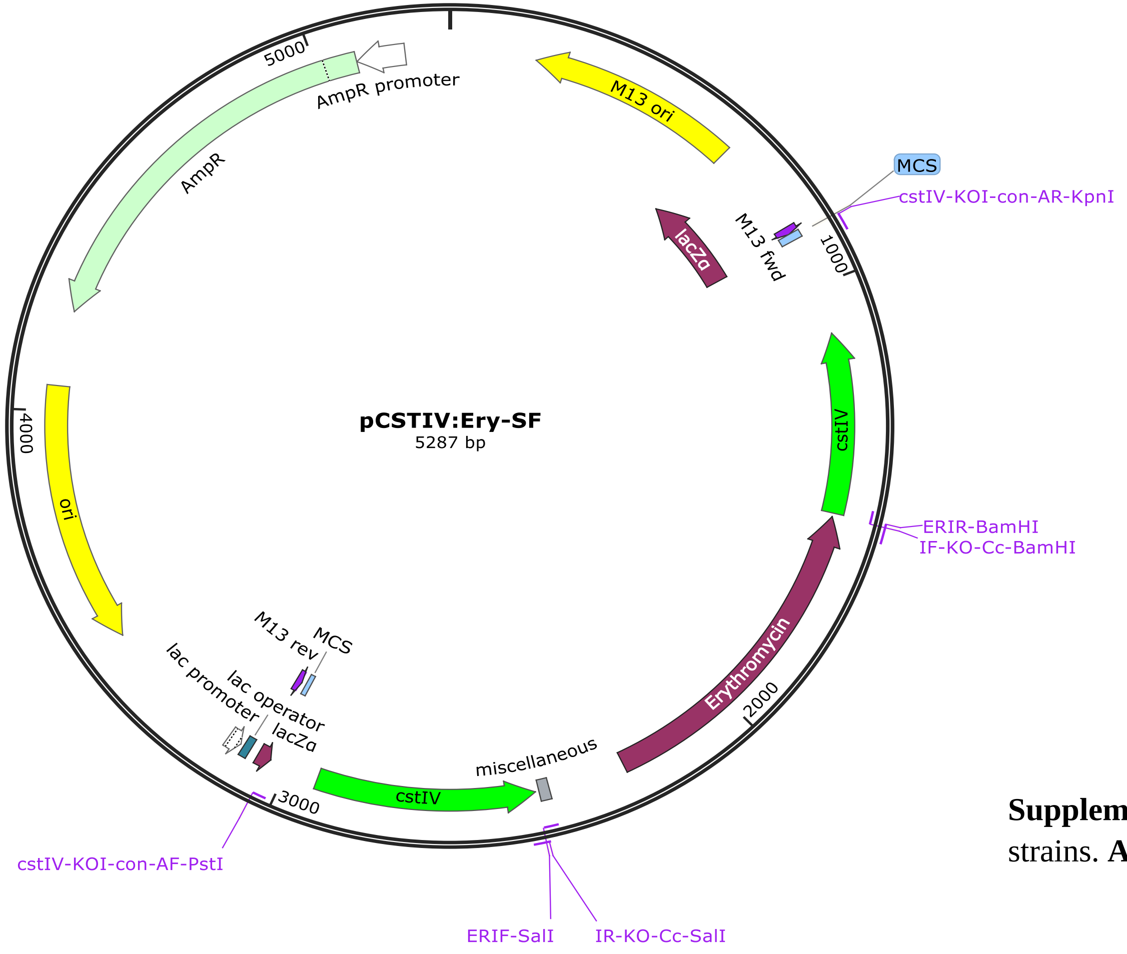


**B**

| Primer name | Sequence |
|---|---|
| ***cstIV* amplification** | |
| **cstIV-KOI-con-AF-PstI** | ATCTGCAGGCCAAAACCACTCACTTAAAAG |
| **cstIV-KOI-con-AR-KpnI** | AGGTACCGCAATCGATACTGATAATTTTAACGCT |
| **Inverse PCR** | |
| **IF-KO-Cc-BamHI** | AGGATCCGCTGGCATGGATTTTGGAGATG |
| **IR-KO-Cc-SalI** | ATAGTCGACTCTTTATAGCCACATGCAACAGC |
| **Erythromycin cassette** | |
| **ERIF-SalI** | ACCGTCGACAGTATAAAACCTTTAAGAACTTTC |
| **ERIR-BamHI** | ACCGGATCCACTTACTTATTAAATAATTTATAGCTAT TG |
| **Mutant verification** | |
| **CcCstIVvF** | AACAGAAATGCTTACTGGCAC |
| **EryCvF2** | ATATTTTCATCCTAAACCTAAAGTGAATAGC |
| **EryCvR1** | TTATTTTCTGTAGTTTTGCATAATTTATGG |

**Supplemental Figure S6.** Plasmids and primers for the generation of *C. coli* 65 and 73 Δ*cstIV* mutant strains. **A;** pCSTIV:Ery-SF plasmid, **B;** primer list

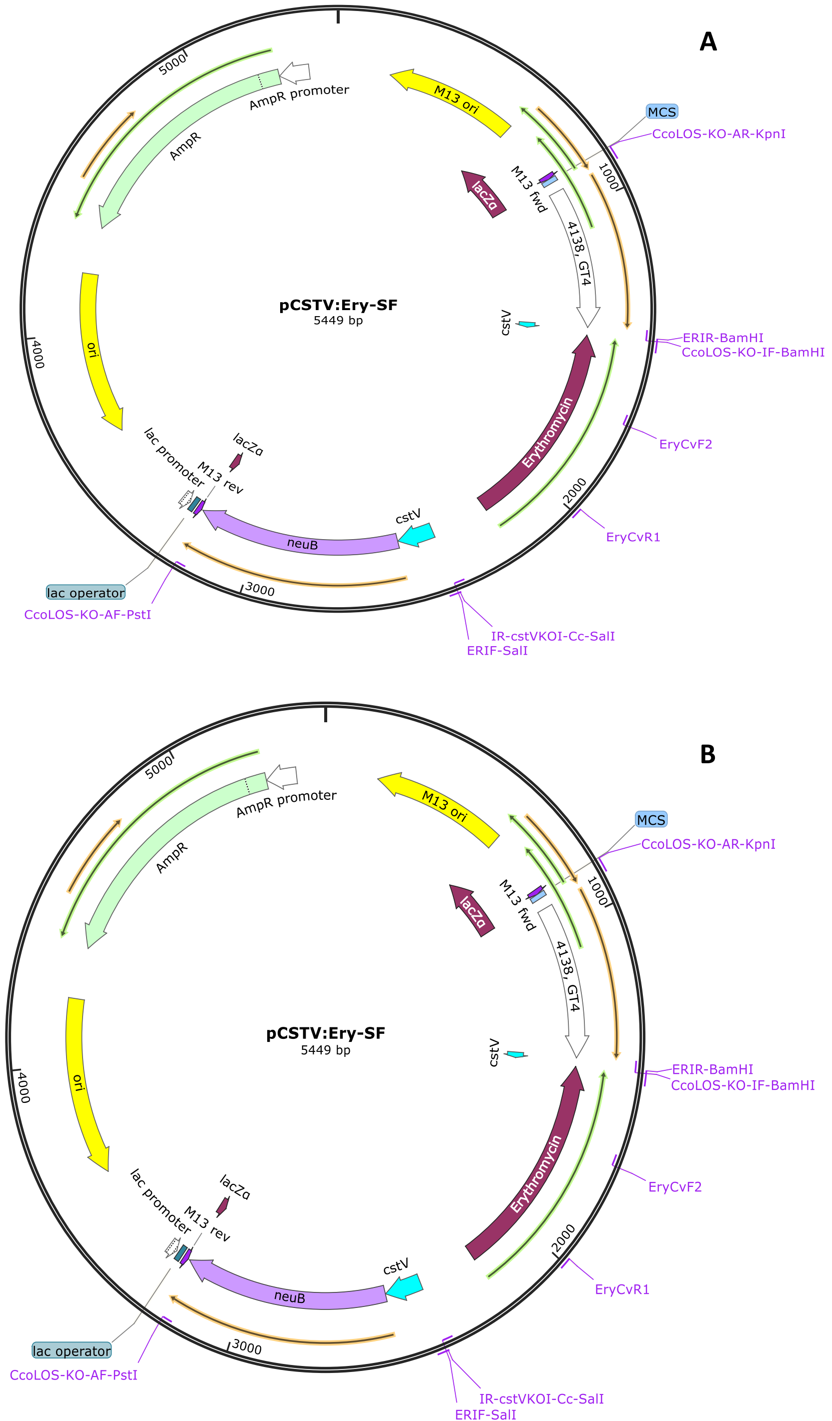


| **Primer name** | **Sequence** |
|---|---|
| **cstV amplification** | |
| **CcoLOS-KO-AF-PstI** | ATCTGCAGCCTAATGCAACCGCACCCAAGC |
| **CcoLOS-KO-AR-KpnI** | AGGTACCATTTATCCCAAAAAGTTGTCTTAAGCGTGG |
| **Inverse PCR** | |
| **IR-cstVKOI-Cc-SalI** | ACCGTCGACCATATCGAAATTATTCTAGAGCTC |
| **CcoLOS-KO-IF-BamHI** | AGGATCCGTTTTCTATCATATCTGTCCTCATAG |
| **Erythromycin cassette** | |
| **ERIF-SalI** | ACCGTCGACAGTATAAAACCTTTAAGAACTTTC |
| **ERIR-BamHI** | ACCGGATCCACTTACTTATTAAATAATTTATAGCTATTG |
| **ERIR-SalI** | ACCGTCGACACTTACTTATTAAATAATTTATAGCTATTG |
| **ERIF-BamHI** | ACCGGATCCAGTATAAAACCTTTAAGAACTTTC |
| **Mutant verification** | |
| **EryCvF1** | CCTCTTATTATAGCCATTTGTTTGC |
| **EryCvF2** | ATATTTTCATCCTAAACCTAAAGTGAATAGC |
| **EryCvR1** | TTATTTTCTGTAGTTTTGCATAATTTATGG |
| **EryCvR2** | GGAGAAAGAGTTTGTGCTAATCTC |
| **Gene expression verification** | |
| **09870cFv** | TAA GTT TTT GGT AGT TTT TGC CTC G |
| **09870cRv** | ATTAGCACTAGATGATACAACCAGTG |
| **09880cFv** | CAACTAAGATCCCATTTATGCCAAG |
| **09880cRv** | TAGAAGGTGGAGAGCTTTCAGG |
| **09890cFv** | ACATAGCCTGTAATCTTACTAAATACG |
| **09890cRv** | ATAGGTTCAGGAGAATGTAATAACTATCC |
| **09910cFv** | GGC TAG TAA GTG CAT AAA TAC TTG C |
| **09910cRv** | CGATCAAAGAAAGTGATTTATCCCAA |
| **09930cFv** | TGCGTGGTAGAGCTAGGATG |
| **09930cRv** | ACAATGAAAGCACTGATGACACTC |

**Supplemental Figure S7.** Plasmids and primers for the generation of *C. coli* 76339 Δ*cstV* mutant strains. **A;** pCSTV:Ery-SF plasmid, **B;** pCSTV:Ery-RF plasmid, C; primer list

A

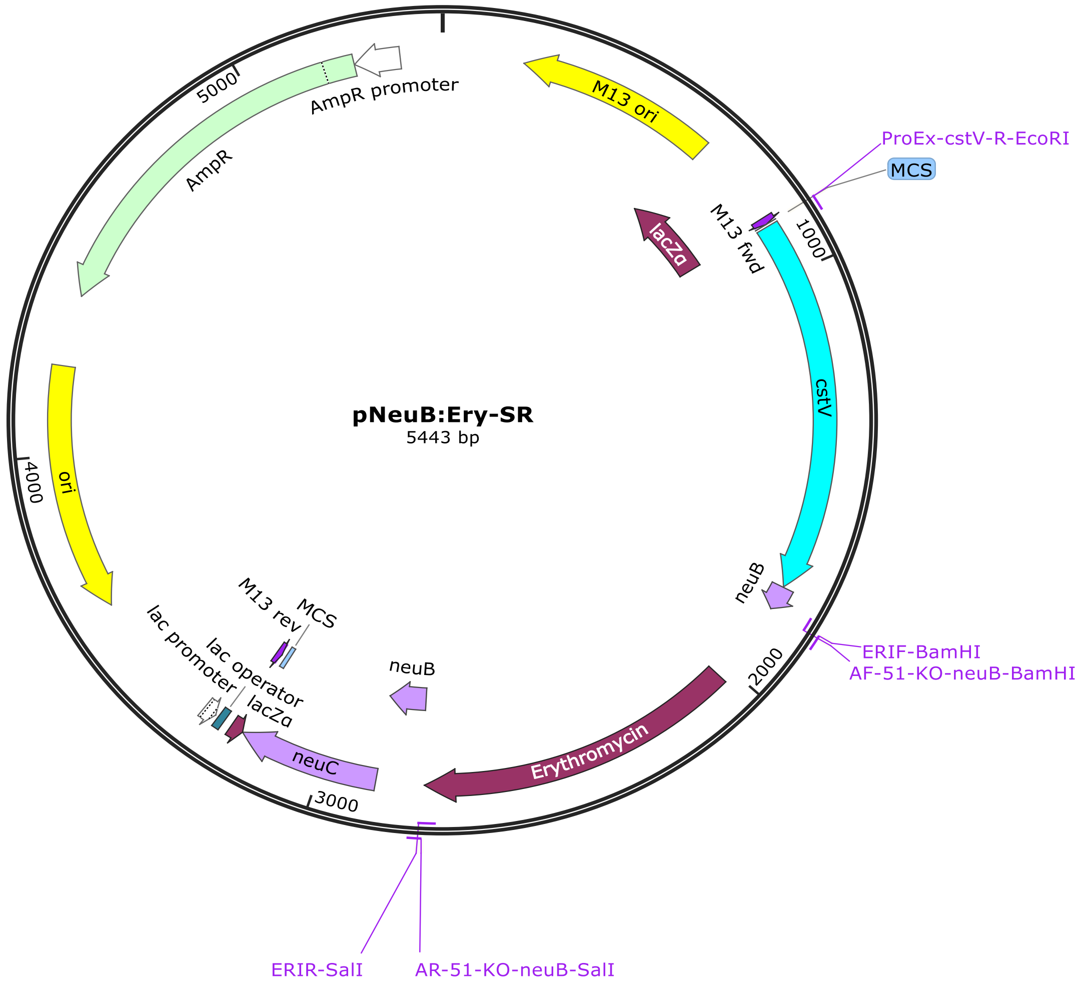


B

| Primer name | Sequence |
|---|---|
| **neuB amplification** | |
| **AF-51-KO-neuB-PstI** | ATCTGCAGCTCTCCACCTTCTATATGTGCTAC |
| **ProEx-cstV-R-EcoRI** | ACTGAATTCATAGAAAACAATGCAGTTGTTGTT |
| **Inverse PCR** | |
| **AR-51-KO-neuB-SalI** | ACCGTCGACGGGACTCGGTGGAATCAGC |
| **AF-51-KO-neuB-BamHI** | AGGATCCCTAAAGGTGGGTTTTCTTGGGATATG |
| **Erythromycin cassette** | |
| **ERIR-SalI** | ACCGTCGACACTTACTTATTAAATAATTTATAGCTATTG |
| **ERIF-BamHI** | ACCGGATCCAGTATAAAACCTTTAAGAACTTTC |
| **Mutant verification** | |
| **EryCvF2** | ATATTTTCATCCTAAACCTAAAGTGAATAGC |
| **EryCvR1** | TTATTTTCTGTAGTTTTGCATAATTTATGG |

**Supplemental Figure S8.** Plasmids and primers for the generation of *C. coli* 76339 Δ*neuB1* mutant strains. A; pNeuB:Ery-SR plasmid, B; primer list

**A**

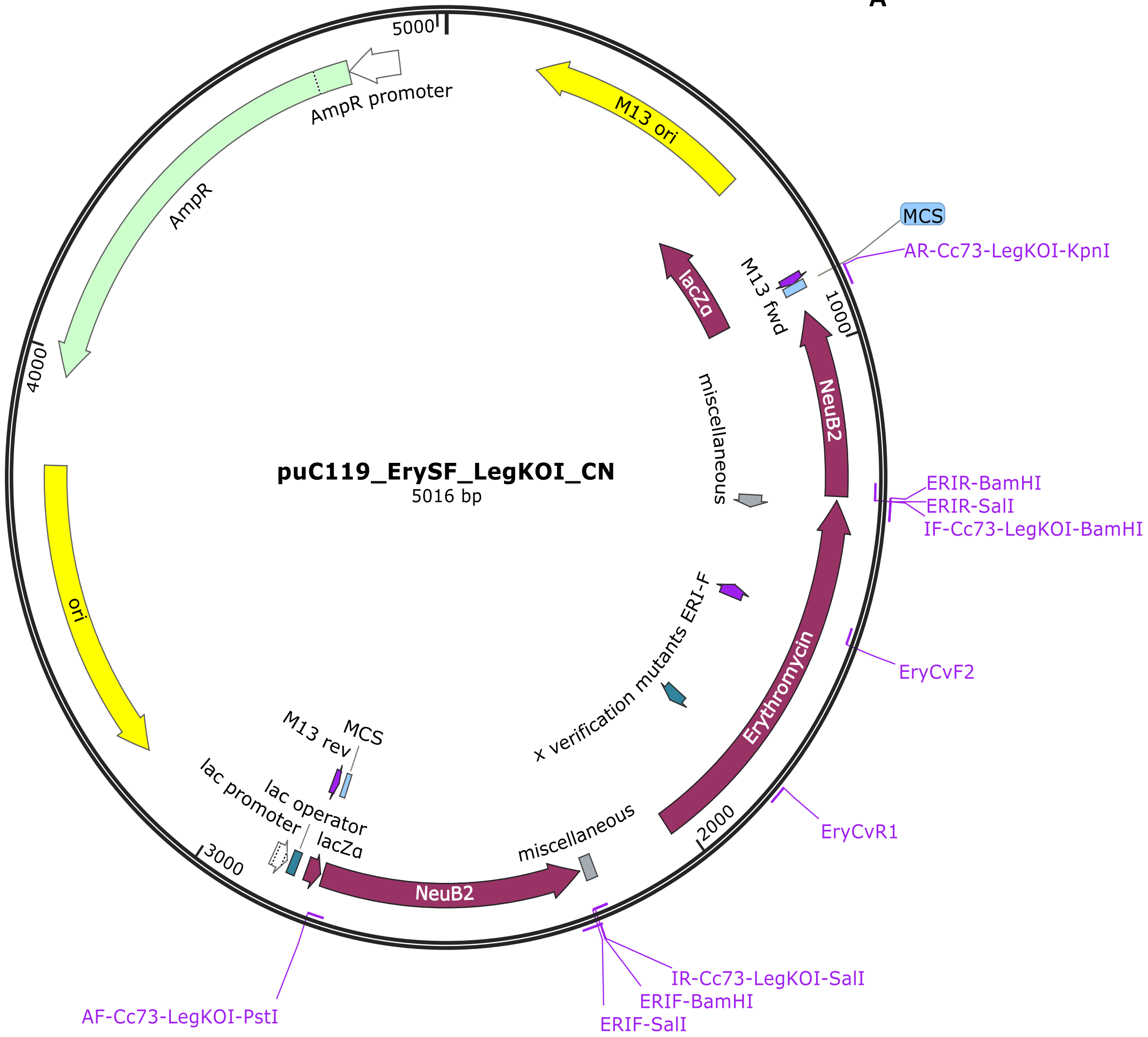


**B**

| **Primer name** | **Sequence** |
|---|---|
| **neuB2 amplification** | |
| **AF-Cc73-LegKOI-PstI** | ATCTGCAGAAAACTCTTATCATCGCAGAAGCAGG |
| **IR-Cc73-LegKOI-SalI** | TATGTCGACAGGGGCAGGGTATTCTGTAGTACAGTG |
| **AR-Cc73-LegKOI-KpnI** | AGGTACCCATTCAGCCCTTGTTGCACTGACTATAC |
| **IF-Cc73-LegKOI-BamHI** | AGGATCCCAGTTGCCTTGGGAGCTTGTGTAATAG |
| **ERIR-SalI** | ACCGTCGACACTTACTTATTAAATAATTTATAGCTAT TG |
| **ERIF-BamHI** | ACCGGATCCAGTATAAAACCTTTAAGAACTTTC |
| **Mutant verification** | |
| **LegKOI-cR** | CATCAATAGACATACACTTGCTACG |

**Supplemental Figure S9.** Plasmids and primers for the generation of *C. coli* 73 Δ*neuB2* mutant strains. A; pLegKOI:Ery-SF plasmid, B; primer list

A

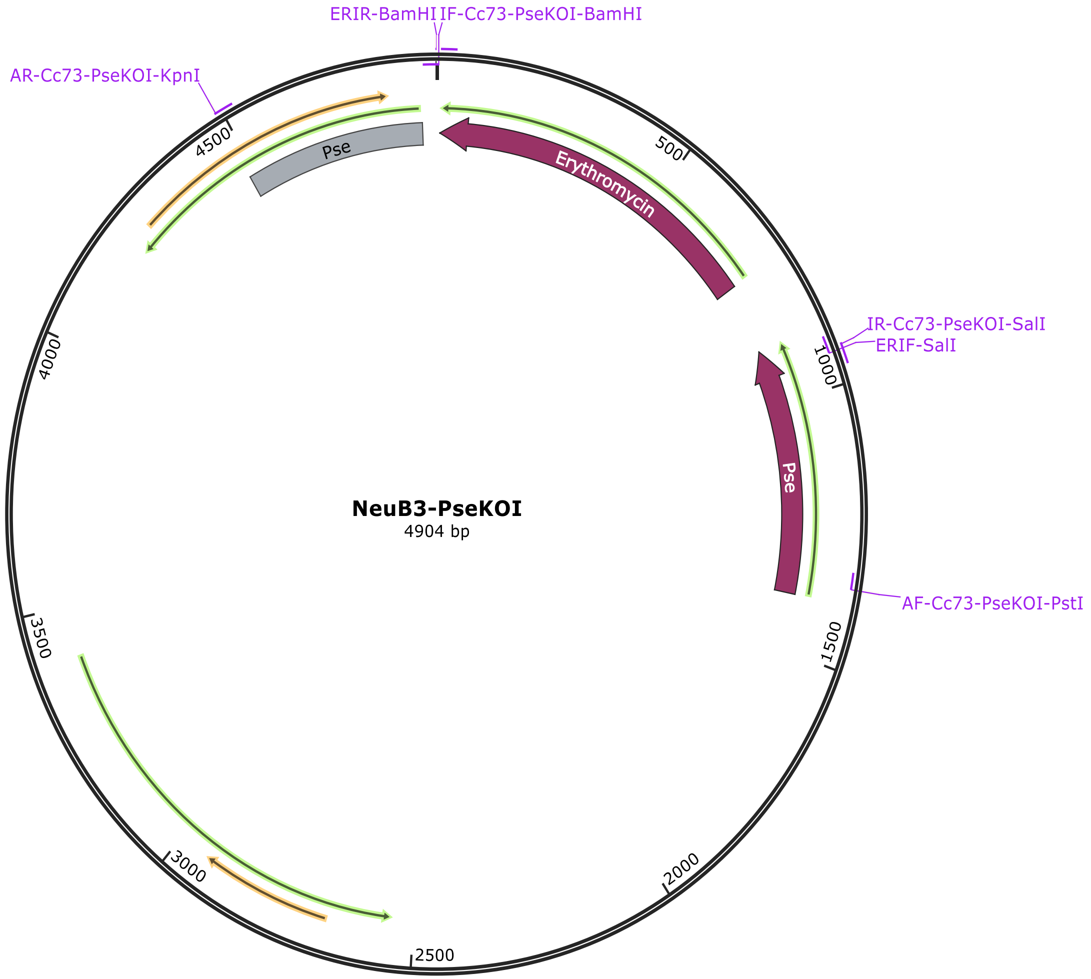


B

| Primer name | Sequence |
|---|---|
| **neuB3 amplification** | |
| **AF-Cc73-PseKOI-PstI** | ATCTGCAGTCACGCAGGAAGTCTTGAGATGGC |
| **AR-Cc73-PseKOI-KpnI** | AGGTACCGGATGTAGTCCAAAAGAAGGACGCACG |
| **IR-Cc73-PseKOI-SalI** | TATGTCGACGTTCTTCTTCTGTAGCAATGCCCGTTG |
| **IF-Cc73-PseKOI-BamHI** | AGGATCCACTTCGGCTTATCCTACTGCCATAG |
| **ERIF-SalI** | ACCGTCGACAGTATAAAACCTTTAAGAACTTTC |
| **ERIR-BamHI** | ACCGGATCCACTTACTTATTAAATAATTTATAGCTATT G |
| **Mutant verification** | |
| **PseKOI-cR** | GAACGACAAGCATCACAAACACC |

**Supplemental Figure S10.** Plasmids and primers for the generation of *C. coli* 73 Δ*neuB3* mutant strains. A; PseKOI plasmid, B; primer list

A

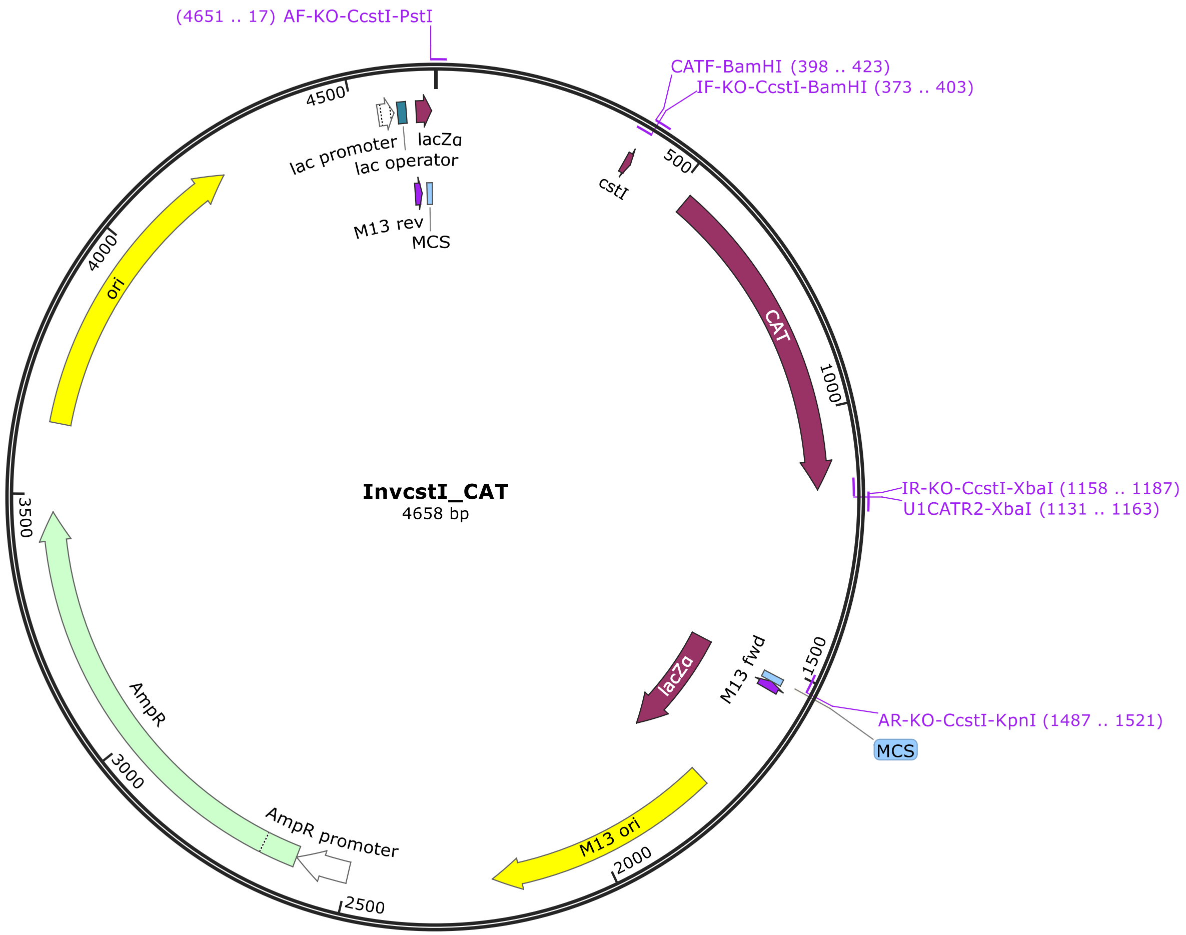


B

| Primer name | Sequence |
|---|---|
| **cstI amplification** | |
| **AF-KO-CcstI-PstI** | ATCTGCAGAGGAAATGGGCCTAGTCTT |
| **AR-KO-CcstI-KpnI** | AGGTACCCACTATGGATTTCATAAGAATTTCTACCT |
| **Inverse PCR** | |
| **IR-KO-CcstI-XbaI** | ATTCTAGATGAATTTGGCTAAGACTGTAGGTG |
| **IF-KO-CcstI-BamHI** | ACCGGATCCGCTATGGCGCACATATAAATACCAG |
| **CAT cassette** | |
| **U1CATR2-XbaI** | ATTCTAGAGGGATTTTATTTATTCAGCAAGTCTTG |
| **CATF-BamHI** | AGGATCCCGGCGGTGTTCCTTTCCAAG |
| **Mutant verification** | |
| **cstI-CvR** | GCATTCTTCTATACAATTTAACTCCC |
| **cstI-CvF** | ACTAAAAAACGGGAGGGAAGC |
| **CAT-CvR** | CTCAGTCCAAATACTCGAAAAGG |
| **CAT-CvF** | TCTATGATACCGTGGACAAGC |

**Supplemental Figure S11.** Plasmids and primers for the generation of *C. coli* 76339 Δ*cstI* mutant strains. A; InvcstI_CAT plasmid, B; primer list

A

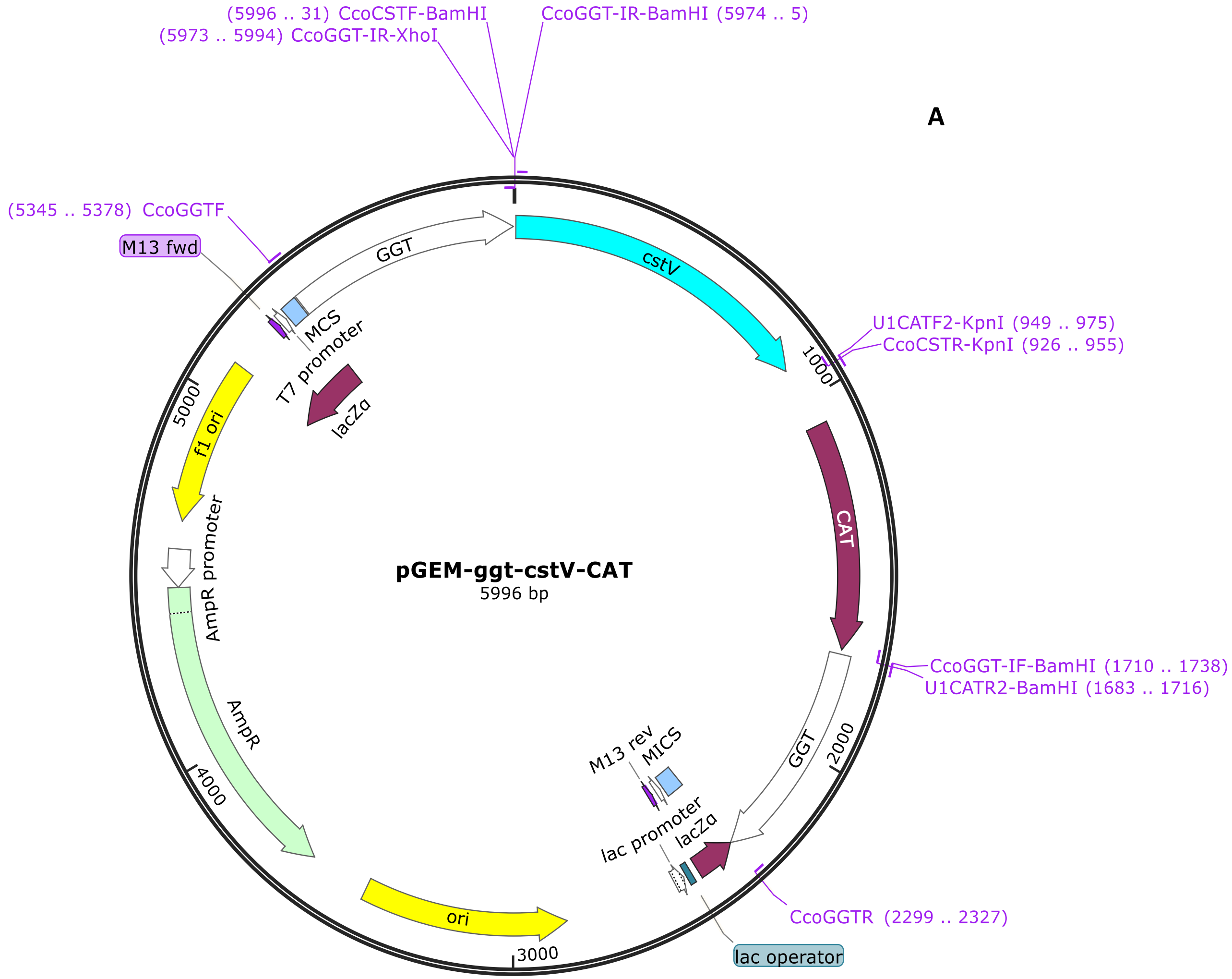


B

| Primer name | Sequence |
|---|---|
| **ggt amplification** | |
| **CcoGGTF** | ATTTAGTTATATTTGTGATTTCAATCACGCTAGG |
| **CcoGGTR** | AAAACTGTTGTGATGATTCTAGAGCCACC |
| **Inverse PCR** | |
| **CcoGGT-IF-BamHI** | AGGATCCTTCTATGTCGCCACCTAGTAGCG |
| **CcoGGT-IR-BamHI** | AGGATCCCAGGGCCTTCTTTGGCGATGAG |
| **cstV amplification** | |
| **CcoCSTR-KpnI** | ACCGGTACCTTCTTGGGATATGGTTAATTTATC |
| **CcoCSTF-BamHI** | AGGATCCAATGATAGAAAACAATGCAGTTGTTG |
| **CAT cassette** | |
| **U1CATR2-KpnI** | AGGTACCGGGATTTTATTTATTCAGCAAGTCTTG |
| **U1CATR2-BamHI** | AGGATCCGGGATTTTATTTATTCAGCAAGTCTTG |

**Supplemental Figure S12.** Plasmids and primers for the generation of *cstV* complemented strain. A; pGEM-ggt-cstV-CAT plasmid, B; primer list
